# Thymidine Phosphorylase Mediates SARS-CoV-2 Spike Protein Enhanced Thrombosis in K18-hACE2^TG^ Mice

**DOI:** 10.1101/2024.02.23.581661

**Authors:** Renat Roytenberg, Hong Yue, Autumn DeHart, Eugene Kim, Fang Bai, Yongick Kim, Krista Denning, Alec Kwei, Quan Zhang, Jiang Liu, X. Long Zheng, Wei Li

## Abstract

COVID-19, caused by SARS-CoV-2, is associated with arterial and venous thrombosis, thereby increasing mortality. SARS-CoV-2 spike protein (SP), a viral envelope structural protein, is implicated in COVID-19-associated thrombosis. However, the underlying mechanisms remain unknown. Thymidine phosphorylase (TYMP), a newly identified prothrombotic protein, is upregulated in the plasma, platelets, and lungs of patients with COVID-19 but its role in COVID-19-associated thrombosis is not defined. In this study, we found that wild-type SARS-CoV-2 SP significantly promoted arterial thrombosis in K18-hACE2^TG^ mice. SP-accelerated thrombosis was attenuated by inhibition or genetic ablation of TYMP. SP increased the expression of TYMP, resulting in the activation of signal transducer and activator of transcription 3 (STAT3) in BEAS-2B cells, a human bronchial epithelial cell line. A siRNA-mediated knockdown of TYMP inhibited SP-enhanced activation of STAT3. Platelets derived from SP-treated K18-hACE2^TG^ mice also showed increased STAT3 activation, which was reduced by TYMP deficiency. Activated STAT3 is known to potentiate glycoprotein VI signaling in platelets. While SP did not influence ADP- or collagen-induced platelet aggregation, it significantly shortened activated partial thromboplastin time and this change was reversed by TYMP knockout. Additionally, platelet factor 4 (PF4) interacts with SP, which also complexes with TYMP. TYMP enhanced the formation of the SP/PF4 complex, which may potentially augment the prothrombotic and procoagulant effects of PF4. We conclude that SP upregulates TYMP expression, and TYMP inhibition or knockout mitigates SP-enhanced thrombosis. These findings indicate that inhibition of TYMP may be a novel therapeutic strategy for COVID-19-associated thrombosis.

**Key Points:**

- SARS-CoV-2 spike protein, thymidine phosphorylase, and platelet factor 4 form a complex that may promote clot formation.
- Inhibiting thymidine phosphorylase attenuates SARS-CoV-2 spike protein-enhanced thrombosis, platelet activation, and coagulation.

## Introduction

Coronavirus disease 2019 (COVID-19), caused by severe acute respiratory syndrome coronavirus 2 (SARS-CoV-2)^1^, has infected more than 770 million people, caused almost 7 million deaths, and left millions of patients needing long-term care, according to WHO data up to the fourth quarter of 2023^2^. The symptoms of COVID-19 vary. However, all patients with severe COVID-19 are associated with a significantly increased risk of arterial thrombosis and/or venous thromboembolism^3–7^, not only in the acute phase but also in the post-COVID period^3^.

Patients with thrombosis have a higher mortality rate than those without thrombosis^8^. Moreover, patients with COVID-19 have increased morbidity from myocardial infarction and stroke that cannot be explained by the severity of the viral load of SARS-CoV-2, as these complications also occur in patients with mild symptoms^9,10^. The mechanism that mediates COVID-19-associated thrombosis remains incompletely elucidated. Various factors, such as platelet hyperactivation^11,12^, endothelial dysfunction^13,14^, immunothrombosis^14^, and/or coagulation system hyperactivation^15^, etc., have been proposed as potential routes of pathology, but no single, universally successful therapeutic strategy targeting these mechanisms has been established. Therefore, a better understanding of the pathogenesis of COVID-19-associated thrombosis and developing novel, mechanism-based, effective therapeutic strategies for such complications remains an active area of research.

SARS-CoV-2 is a single-strained RNA virus, encoding 29 proteins. While no data show the correlation between the 16 non-structural proteins and the 9 accessory proteins with thrombosis, structural proteins such as nucleocapsid (N), envelope (E), and especially spike protein (SP) have been implicated in enhancing thrombosis^16–18^. Through binding to heparan sulfate, SP increases thrombin activity, leading to direct thrombosis in a zebrafish model in vivo^19^. SP has also been shown to directly activate platelets^20^, which remains to be controversial^15,21^.

SP is detectable in plasma up to a year of post-infection in patients experiencing post-acute sequelae of COVID-19^22^, which coincides with an increased risk of thrombosis in this patient population^3^. This observation aligns with some clinical manifestations of patients who received SARS-CoV-2 vaccination. All major COVID-19 vaccines lead to increased SP expression and are associated with vaccine-induced immune thrombotic thrombocytopenia (VITT)^23^. SP has also been detected in the plasma of patients vaccinated against COVID-19^24,25^ and the participation of SP has been suggested to play a role in VITT^23,26^. However, the initiating mechanism of VITT remains unclear.

Thymidine phosphorylase (TYMP), known as platelet-derived endothelial cell growth factor, is highly expressed in platelets. Recent studies have demonstrated that TYMP participates in platelet signaling and enhances thrombosis^27,28^. Several studies have demonstrated that TYMP is significantly increased in various cells or tissues of COVID-19 patients, especially in the acute phase^29–33^. Using a database provided by Olink and Massachusetts General Hospital (MGH)^29^, we found that TYMP is significantly increased in COVID-19 patients, and the increase of TYMP is positively associated with the severity of COVID-19 and thrombotic events^31^. An in silico-based drug screening study also suggested that TYMP could be a target for treating COVID-19 and tipiracil, a selective TYMP inhibitor, inhibits viral amplification and SP production^34^. Based on these data, we hypothesize that TYMP is a key factor for SP-enhanced thrombosis and inflammation. Targeting TYMP could be a potentially new therapeutic strategy for COVID-19-associated thrombosis.

## Methods

Detailed methods can be found in the online Supplemental Methods and Materials. All animal studies have been approved by the Institutional Animal Care and Use Committee (IACUC) of Marshall University (Protocol #1033528, PI: Wei Li). Human study protocol was approved by the Internal Review Board of the University of Kansas Medical Center, Kansas City, KS (STUDY00148313).

To generate a reliable source of SP for this project, we received plasmids encoding WT SARS-CoV-2 SP (Wuhan-Hu-1) and its receptor binding domain (RBD) from Addgene under a Material Transfer Agreement. The SP- and RBD-encoding plasmids, along with the empty vector pCDNA3.1 (p3.1), were transiently transfected into COS-7 cells. Crude cell lysates were prepared in PBS under sterile conditions 36 hours later. The concentration of SP or RBD was determined by western blot, with a commercially purchased SP S1 subunit serving as a quantitative control (**Fig. S1**), and we estimated that approximately 3% of the total cell lysate was SP or RBD by mass.

Since wild type C57BL/6J (WT) mice do not respond well to SARS-CoV-2 SP, we purchased K18-hACE2^TG^ mice from The Jackson Laboratory. These mice were then crossed with *Tymp^-/-^* mice^27^ to generate a new K18-hACE2^TG^/*Tymp^-/-^* mouse strain. Mice aged 8-16 weeks were used. They were treated with SP- or p3.1-containing cell lysates (500 µg/mouse) via intraperitoneal injection and then used for subsequent studies three days later. Some mice that received SP were also simultaneously administered tipiracil at a dose of 1 mg/kg/day via gavage feeding for three days. These mice were subjected to the 7.5% FeCl_3_-induced carotid artery thrombosis model^35,36^ for in vivo thrombosis assessment, or whole blood was drawn for platelet function assays in vitro, measuring activated partial thromboplastin time (aPTT), and platelet biochemistry assays.

To clarify the pathophysiological consequences underlying SP treatment, we treated BEAS-2B cells, a human bronchial epithelial cell line (ATCC), with COS-7 lysates containing SP, RBD, or p3.1. Subsequently, cells were harvested for western blot assay of targeting molecules indicated in the Results section. For an in-depth mechanistic study, we conducted co-transfection of plasmids encoding human TYMP and human platelet factor 4 (PF4) along with SP. Cell lysates from these experiments were used for co-immunoprecipitation (co-IP) or Blue Native PAGE assays^37^ to determine the interactions among TYMP, PF4, and SP. Additionally, COS-7 cells co-transfected with SP and PF4 were subjected to immunocytofluorescence to assess the colocalization of PF4 and SP. Furthermore, human plasma isolated from COVID-19 patients and a pooled healthy donor plasma isolated in 2008 were used for co-IP assays to investigate the association between SP and PF4. We also employed AlphaFold2 in silico structure analysis to predict the binding interface between TYMP and SP as well as SP and PF4 through the ChimeraX interface^38,39^.

To further study the potential effect of the SP and PF4 on platelet function, we treated whole blood isolated from K18-hACE2^TG^ and K18-hACE2^TG^/*Tymp^-/-^* mice with COS-7 lysates containing murine PF4 (mPF4) alone or a combination of mPF4 and SP for one hour at 37 °C. Subsequently, they are perfused through a collagen-coated Vena8 Fluoro+ 8-channel microfluidic flow chamber at a constant flow rate of 60 µl/min for three minutes. The chamber was then washed with PBS at the same flow rate for another three minutes and fluorescence areas taken at five predetermined marker sites were quantified using ImageJ.

The data were analyzed for normality and equal variance as a justification for using the 2-tailed Student’s *t* test, One-way ANOVA with Dunnett’s multiple comparisons test, and Two-way ANOVA with Dunnett’s multiple comparisons test where applicable. The logrank test using the Kaplan-Meier survival curve was used to analyze the flow cessation data. GraphPad Prism Version 10.1.2 (GraphPad, USA) was used for data analysis. Data were presented as Mean ± SE and a *p* ≤ 0.05 was considered statistically significant.

## Results

### SARS-CoV-2 spike protein enhances thrombosis in K18-hACE2^TG^ mice, which is inhibited by genetic knockout of TYMP or by tipiracil

Previous studies have indicated that SARS-CoV-2 SP does not bind well to mouse angiotensin converting enzyme 2 (ACE2), which serves as the receptor mediating SARS-CoV-2 entry into host cells^40,41^. We thus employed the use of K18-hACE2^TG^ mice, a widely used animal model for studying SARS-CoV-2 invasion and therapeutic strategies^42^. This transgenic mouse model expresses human ACE2 in most epithelial cells driven by the human keratin 18 promoter (K18)^43^. Male K18-hACE2^TG^ mice without any treatment were subjected to the 7.5% FeCl_3_-induced carotid artery injury thrombosis model. Unexpectedly, these mice showed an antithrombotic phenotype and significantly prolonged times to forming occlusive thrombi compared to WT mice (data not shown). Subsequently, we treated K18-hACE2^TG^ mice with 500 µg p3.1- or SP-containing lysates by intraperitoneal injection. The in vivo thrombosis study was conducted three days later. As shown in **Fig. 1A** and **1B**, as well as **Videos 1 and 2**, p3.1 did not significantly affect thrombosis in the K18-hACE2^TG^ mice compared to normal K18-hACE2^TG^ mice (data not shown); however, SP treatment significantly shortened blood flow cessation times.

**Figure 1:**
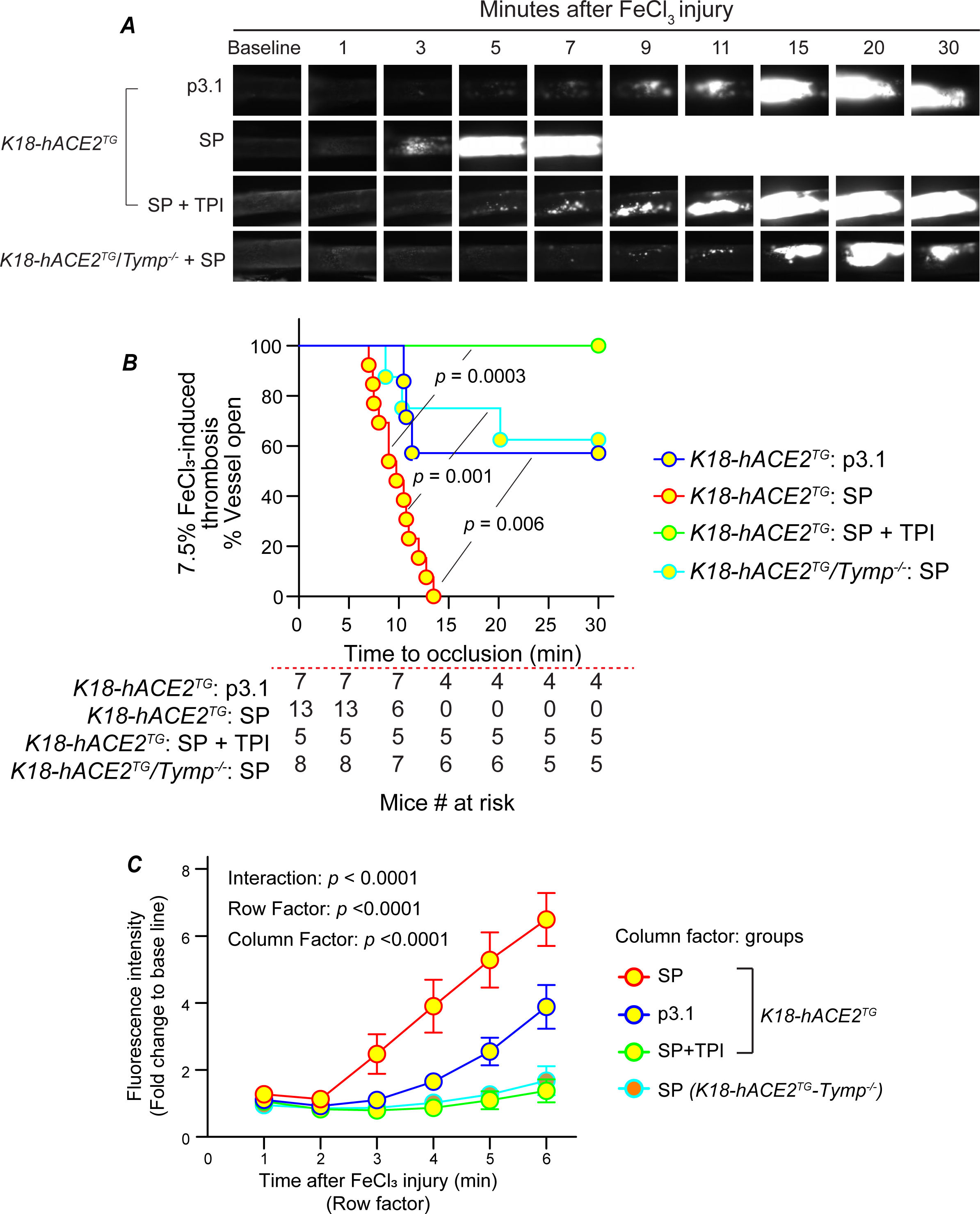
TYMP deficiency or inhibition attenuates SARS-CoV-2 spike protein-enhanced thrombosis in K18-hACE2^TG^ mice. K18-hACE2^TG^ and K18-hACE2^TG^/*Tymp^-/-^*mice were injected intraperitoneally with 500 μg SARS-CoV-2 spike protein-containing (SP) or control (p3.1) COS-7 lysate. Tipiracil hydrochloride (TPI) was administered at 1 mg/kg once every 24 hours to one group of K18-hACE2^TG^ mice treated with SP. The 7.5% FeCl_3_-induced carotid artery thrombosis model was performed 72 hours after SP or control injection. ***A***. Still images taken of murine carotid arteries during the assay. ***B***. The survival plot of time to flow cessation was analyzed. Groups were compared using the logrank test. ***C***. Fluorescence intensity was measured for the first six minutes after injury and was analyzed via a two-way ANOVA.

Since both TYMP expression and thrombotic events are correlated with COVID-19 disease severity^31,47^, and TYMP expression is highly increased in platelets^32^, plasma^29^, and lungs^30^ of COVID-19 patients, we hypothesized that TYMP plays a role in SP-enhanced thrombosis. We validated this hypothesis by treating K18-hACE2^TG^/*Tymp^-/-^*mice with SP. As shown in Fig. 1A and 1B (cyan curve) as well as **video 3**, TYMP deficiency significantly attenuated SP-enhanced thrombosis. Simultaneously treating SP-treated K18-hACE2^TG^ mice with tipiracil at a dose of 1 mg/kg/day completely diminished SP-enhanced thrombosis (Fig. 1B, green curve, and **video 4**). To further determine how SP affects thrombi progressions, one still image at each minute interval was randomly selected for the first 6 minutes of the assay, and fluorescence intensity was analyzed using ImageJ. Data were presented as fold change compared to baseline (before FeCl_3_ injury) in each individual mouse. As shown in **Fig. 1C**, a steep slope was observed in the SP-treated K18-hACE2^TG^ mice, suggesting accelerated thrombi formations compared to the mice in other groups.

### SARS-CoV-2 spike protein treatment does not influence agonist-induced platelet activation but enhances coagulation

To clarify the mechanism underlying SP-enhanced thrombosis, we investigated whether SP treatment alters platelet aggregation using platelet-rich plasma (PRP). PRP was prepared from SP-treated K18-hACE2^TG^ or K18-hACE2^TG^/*Tymp^-/-^* mice, as well as p3.1-treated K18-hACE2^TG^ mice. These treatments did not affect platelet counts (**Fig. S2**). As shown in **Fig. 2A** and **2B**, we did not observe a difference in collagen- or ADP-induced platelet aggregation.

**Figure 2:**
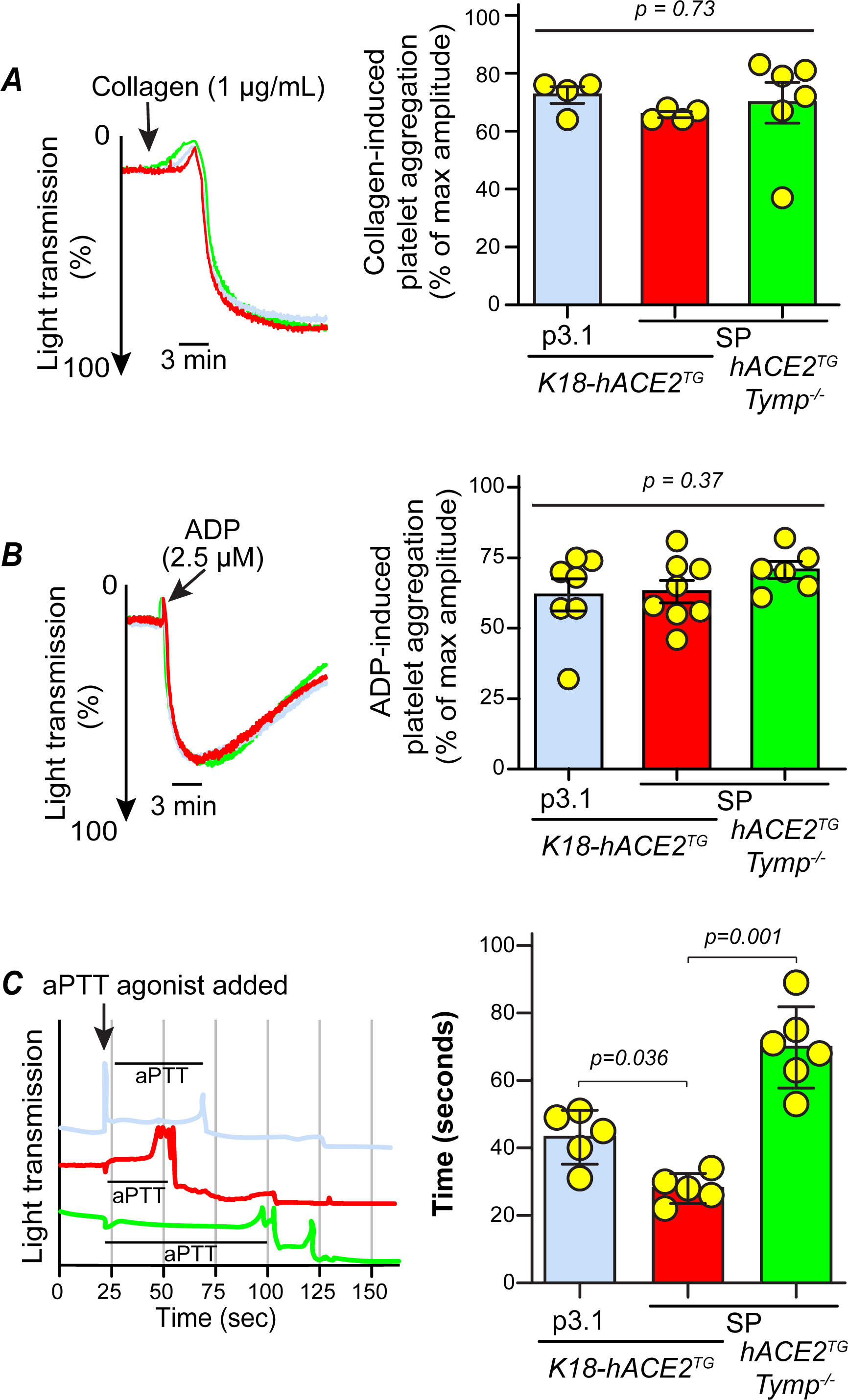
SARS-CoV-2 spike protein does not enhance ADP- or collagen-induced platelet aggregation, but significantly shortened activated partial thromboplastin time. Platelet-rich plasma (PRP) and platelet-poor plasma (PPP) were harvested from K18-hACE2^TG^ and K18-hACE2^TG^/*Tymp^-/-^* 72 hours after treatment with SP- or p3.1-lysate. The maximal light transmitted after agonist induction was used for statistical analyses. *A.* 1 μg/mL collagen- and *B.* 2.5 μM ADP-induced platelet activation. *C.* Activated partial thromboplastin time (aPTT) was measured with the R2 Diagnostics aPTT kit using PPP. Data were analyzed with one-way ANOVA with Dunnett’s multiple comparisons test.

The stability of a thrombus in a high shear arterial vessel (e.g., carotid artery) partially depends on the activity of the intrinsic pathway of the coagulation cascade^44^. Consequently, we conducted an aPTT assay using platelet-poor plasma (PPP) isolated from mice as mentioned above. As depicted in **Fig. 2C**, aPTT was significantly shortened in the SP-treated K18-hACE2^TG^ mice compared to the p3.1-treated K18-hACE2^TG^ mice or SP-treated K18-hACE2^TG^/*Tymp^-/-^*mice.

### SARS-CoV-2 spike protein increases TYMP expression and TYMP deficiency attenuates spike protein-promoted STAT3 activation

The mechanism underlying the increase of TYMP in the COVID-19 milieu is not clear. TYMP is expressed in bronchi and certain alveolar cells (**Fig. 3A**, image is from The Human Protein Atlas). By double immunofluorescence staining of human lung tissue for TYMP and prosurfactant protein C (PSP-C), a marker for type II alveolar cells, we confirmed that these TYMP expressing cells are type II alveolar cells (**Fig. 3B**). Both bronchial epithelial cells and type II alveolar cells express ACE2, which mediates SARS-CoV-2 entry^45,46^. To test if SARS-CoV-2 SP alters the protein expression of TYMP, we treated BEAS-2B cells with conditioned COS-7 lysates. As shown in **Fig. 3C**, both SP- and RBD-containing COS-7 lysate treatments significantly upregulated the expression of TYMP, with an approximately three-fold increase caused by SP.

**Figure 3:**
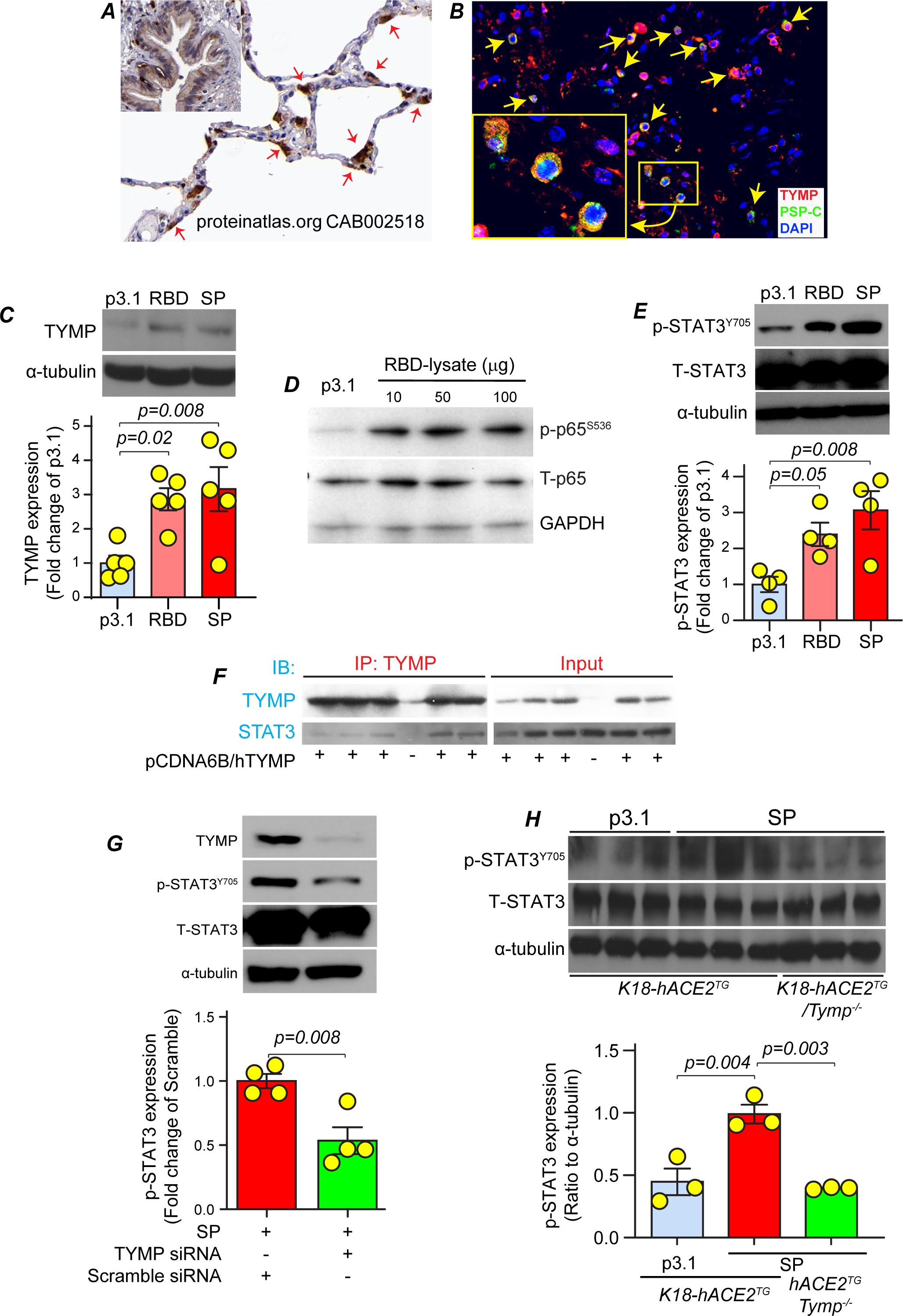
SARS-CoV-2 spike protein enhances TYMP expression and STAT3 activation in BEAS-2B cells and the inhibition of TYMP reduces STAT3 activation. ***A.*** Normal human lung was immunohistochemically stained for TYMP. The image was adopted from “The Human Protein Atlas” (CAB002518). Brown indicates positive staining. ***B.*** Normal human lung tissue was double stained for TYMP and prosurfactant protein C (PSP-C). Nuclei were stained with DAPI. BEAS-2B cells were treated with 50 μg/mL SARS-CoV-2 spike protein (SP)-, RBD-, or p3.1-containing COS-7 lysate and protein expression of (***C***) TYMP, (***D***) p-p65, and (***E***) p-STAT3 were examined by western blot 24 hours later. ***F**.* COS-7 cells were transfected with or without pCDNA6B-hTYMP plasmid and then cell lysates were used for immunoprecipitation of TYMP followed by immunoblotting for STAT3 and TYMP. ***G**.* BEAS-2B cells were transfected with 5 nM siRNA targeting TYMP or scramble siRNA for 48 hours. The cells were then treated with 50 μg/mL SP-containing COS-7 lysate for 24 hours and the expression of relevant proteins was examined by western blot. ***H**.* Washed platelets were prepared from p3.1-treated K18-hACE2^TG^ as well as SP-treated K18-hACE2^TG^ and K18-hACE2^TG^/*Tymp^-/-^*mice, and expression of the Y705-phosphorylated and total STAT3 were determined by western blot. Alpha-tubulin was blotted as a control. Data in *C*, *E*, and *H* were analyzed by one-way ANOVA with Dunnett’s multiple comparisons test and *G* were analyzed by unpaired Student’s *t*-test.

Both nuclear factor kappa-light-chain-enhancer of activated B cells (NF-κB) and signal transducer and activator of transcription 3 (STAT3) signals rapidly respond to stimuli and have been implicated in COVID-19-associated thrombosis and inflammation^47–49^. NF-κB is known to upregulate TYMP expression^50^. Therefore, we examined the potential correlation among NF-κB, STAT3, and TYMP. As shown in **Fig. 3D and E**, SP or RBD treatment significantly increased the activation of both NF-κB and STAT3 in BEAS-2B cells. STAT3 contains an SH2 domain, which houses Tyr705, a residue necessary for activation and subsequent dimerization^51^. TYMP has previously been shown to bind Src Homology 2/3 (SH2, SH3) domains^28^. Therefore, we examined their interaction and found STAT3 was co-immunoprecipitated with TYMP, suggesting a direct interaction between these two proteins (**Fig. 3F**). A previous study has indicated that TYMP increases serum stimulated STAT3 activation in vascular smooth muscle cells^52^. To clarify the relation between TYMP and STAT3, we knocked down TYMP expression using siRNA and then examined SP-enhanced STAT3 activation. As shown in **Fig. 3G**, TYMP knockdown almost halved STAT3 activation in SP-treated BEAS-2B cells. These data suggest a potential signaling cascade involving NF-κB, TYMP, and STAT3.

STAT3 may potentiate glycoprotein VI (GPVI)-mediated platelet activation^53,54^. Therefore, we examined STAT3 activity in platelets harvested from the SP-treated K18-hACE2^TG^ or K18-hACE2^TG^/*Tymp^-/-^* mice, or K18-hACE2^TG^ treated with p3.1. As shown in **Fig. 3H**, SP significantly increased STAT3 activation in K18-hACE2^TG^ platelets compared to platelets harvested from p3.1-treated K18-hACE2^TG^ mice. TYMP deficiency significantly reduced SP-enhanced STAT3 activation in platelets. Notably, TYMP deficiency did not affect Janus kinase 2 (JAK2) activity (**Fig. S3**), suggesting a TYMP-mediated, SP-enhanced STAT3 activation underlying a mechanism independent of canonical JAK-STAT signaling.

### SARS-CoV-2 spike protein forms a complex with TYMP as well as platelet factor 4

In recent reviews from our lab and others, it has been proposed that SARS-CoV-2 SP potentially interacts with PF4^23,26^. As mentioned above, SP has been found in the plasma of COVID-19 patients and in patients inoculated with COVID-19 vaccines. By co-transfection of plasmids encoding SP or human PF4 (hPF4), followed by immunocytochemistry, we observed the colocalization of SP and hPF4 in some cellular organelles (**Fig. 4A**). To validate this finding, we conducted a co-IP assay using COS-7 cells co-transfected with SP and hPF4 followed by immunoblotting of the eluates. As shown in **Fig. 4B**, pulling down hPF4 also resulted in the pull-down of SP, and vice versa. Additionally, a blue native PAGE assay also demonstrated that SP and PF4 form a complex (**Fig. 4C**). Subsequently, we examined whether TYMP can form a complex with SP and PF4. Using the same strategy and expressing TYMP and SP or PF4 in COS-7 cells, followed by a co-IP assay, we found that TYMP was associated with SP (**Fig. 4D**), but not PF4 (data not shown).

**Figure 4:**
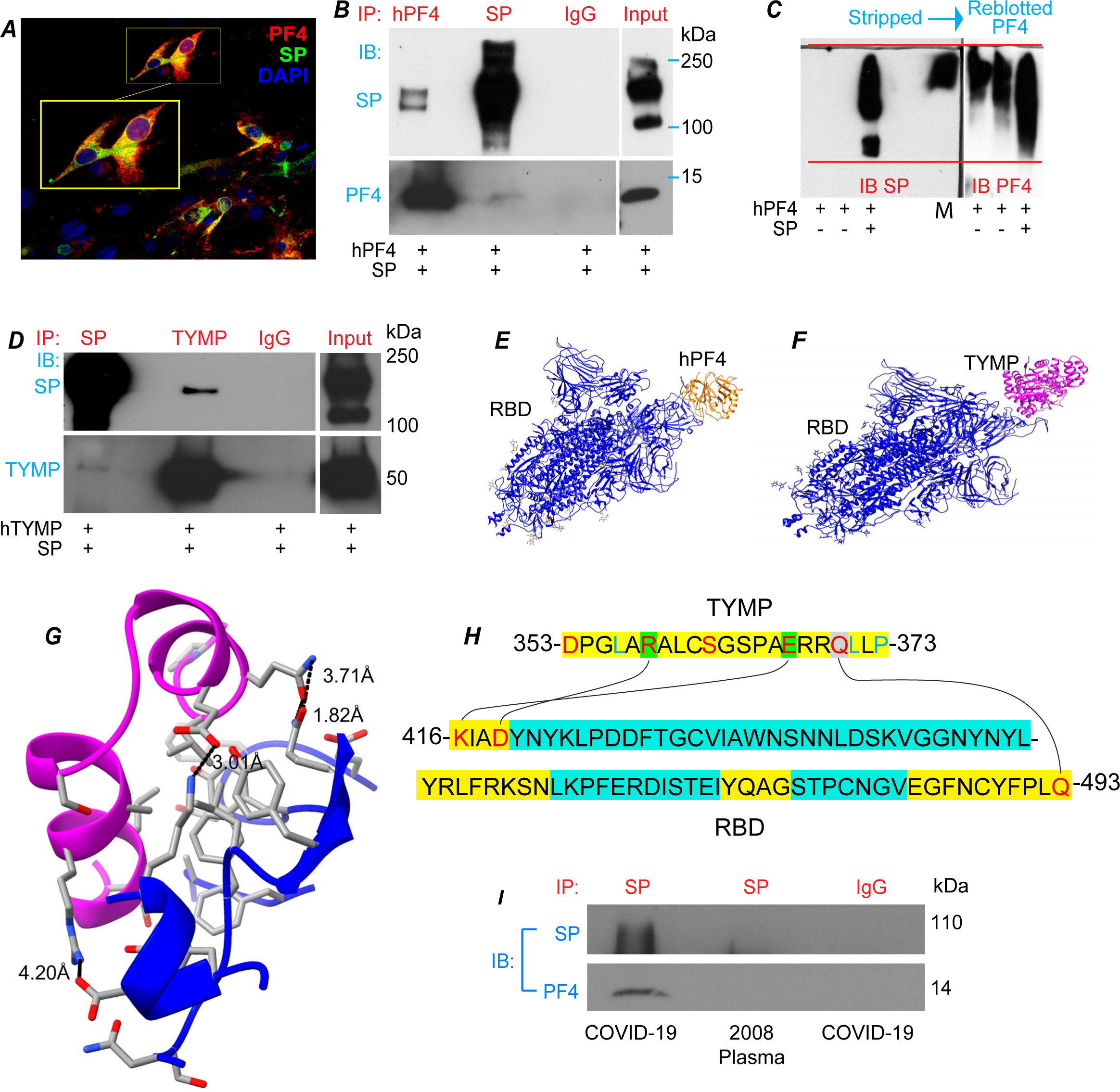
SARS-CoV-2 spike protein forms a complex with TYMP or platelet factor 4. COS-7 cells were co-transfected with plasmids encoding human platelet factor 4 (hPF4) and SARS-CoV-2 SP. ***A.*** The expression of hPF4 and SP was examined by immunocytofluorescence. ***B.*** Cell lysates were prepared and used for IP hPF4 and SP, and then IB for SP and hPF4. IP with normal mouse IgG served as an isotype control. ***C.*** COS-7 cells receiving hPF4 and SP gene transfections were subjected to a Blue Native Polyacrylamide Gel Electrophoresis (BN-PAGE) assay. The membrane was blotted for SP, stripped, and then reblotted for hPF4. ***D.*** COS-7 cells were co-transfected with plasmids encoding SARS-CoV-2 SP and human TYMP, and the resulting cell lysate was used for IP SP and TYMP, and then IB for TYMP and SP. IP with normal mouse IgG served as an isotype control. Panels of B, C, and D represent at least two biological repeats. AlphaFold2-based binding prediction analyses were conducted to determine the binding between (***E***) SP and hPF4, as well as (***F***) SP and hTYMP. ***G.*** Predicted amino acid binding sites between SP and TYMP. Å=angstrom. ***H***. Predicted binding amino acids form salt bridges between peptide sequences 353-373 on TYMP and 416-493 on SP. ***I.*** Human plasma isolated from a severe COVID-19 patient and pooled plasma from healthy subjects isolated in 2008 were used for IP SP, and then immunoblotted for SP and PF4. IP with rabbit IgG served as an isotype control.

To further understand how TYMP binds to SP as well as how SP binds to PF4, we conducted an in-silico binding assay and predicted potential binding sites between SARS-CoV-2 SP S1 subunit, which contains the N terminal domain (NTD) and the RBD, and TYMP or PF4. There was no potential binding observed between the S1 NTD and either TYMP or PF4 (data not shown). There was no collision observed between RBD and PF4 (**Fig. 4E**), however, as they were too close to each other, a potential binding may need some conformational changes in these two proteins. There was no collision between RBD and TYMP and they arranged very well, indicating a strong potential binding between these two proteins (**Fig. 4F**). As shown in **Fig. 4G**, the binding of TYMP and RBD is potentially mediated by a DPGLARALCSGSPAERRQLLP (353D – 373P) peptide in TYMP and a (416K – 493Q) peptide in RBD. The 416K and 419D in RBD potentially interact with 367E and 358R in TYMP via a salt bridge, respectively (**Fig. 4H**). The 493Q in RBD may also form a hydrogen-bond with 370Q in TYMP.

To determine if these observations could be translated to the COVID-19 milieu, we conducted a co-IP assay using plasma harvested from a severe COVID-19 patient and a plasma pool isolated in 2008 as a control. While the SP titer in plasma is likely below the sensitivity of western blot detection (**Fig. S4**), we successfully pulled down SP and found the presence of PF4 in the complex (**Fig. 4I**). Taken together, these data strongly suggest that TYMP and SP, as well as SP and hPF4, can directly form a complex, which may play some role(s) in the COVID-19 milieu.

### TYMP enhances the expression of SARS-CoV-2 spike protein and platelet factor 4 and increases the association of SP/PF4

The mechanism underlying SP-shortened aPTT, as well as the role of TYMP, is not clear. We hypothesized that the binding of SP and TYMP leads to SP confirmational changes, subsequently enhancing the binding between SP and PF4. To test this hypothesis, we co-transfected plasmids encoding these three proteins and maintained the SP/PF4 ratio at one-to-one in mass. Three doses of TYMP were tested, and empty vector pCDNA3.1 was used to balance the total amount of DNA transfected. As depicted in **Fig. S5**, although there was no significant difference, co-transfection of TYMP tended to dose-dependently enhance the expression of both SP and PF4. There was no difference in PF4 pulled down at each condition; however, along with the increase of TYMP dosage, the amount of SP pulled down by PF4 was increased (**Fig. 5A**). These data suggest that TYMP dose-dependently enhances the association of SP/PF4.

**Figure 5:**
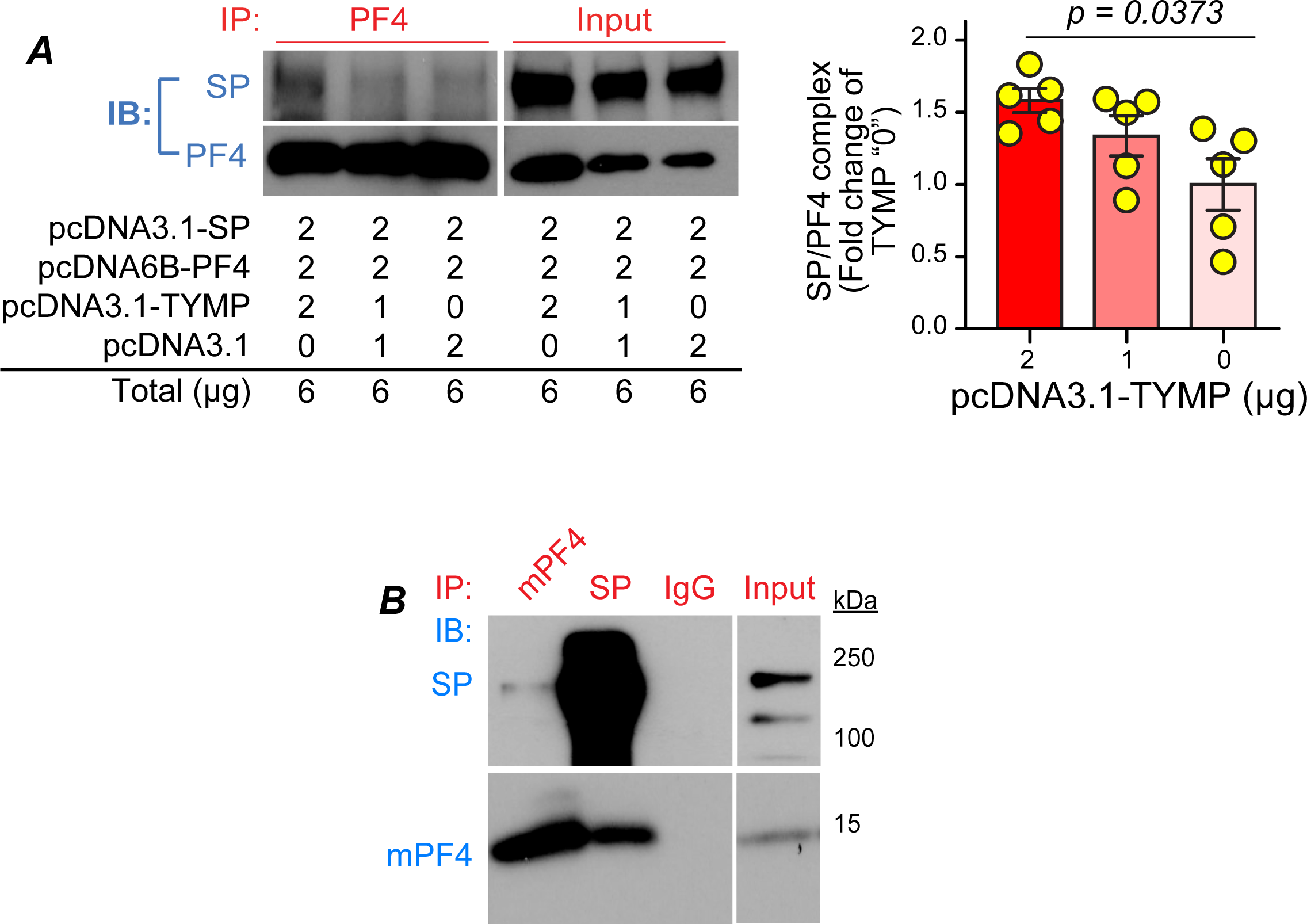
TYMP enhances the complex formation of SARS-CoV-2 spike protein and platelet factor 4. ***A.*** COS-7 cells were co-transfected with plasmids encoding human platelet factor 4 (hPF4), SARS-CoV-2 SP, and human TYMP in various concentrations. The cell lysates were then used for IP PF4 and then blotted for SP and hPF4. Data were analyzed with one-way ANOVA. ***B.*** COS-7 cells were co-transfected with plasmids encoding murine platelet factor 4 (mPF4) and SARS-CoV-2 SP, and the resulting cell lysate was used for co-IP assay of mPF4 and SP. IP with normal mouse IgG served as an isotype control.

To test the functional implication of the SP/PF4 complex, we first examined if SP also binds to mPF4. As shown in **Fig. 5B**, by co-transfection of plasmids encoding SP and mPF4 into COS-7 cells, following a co-IP assay, we found that SP binds to mPF4 too. However, treating WT mice with SP did not affect thrombosis in vivo (**Fig. S6**), suggesting that a SP-hACE2 associated effect is necessary for SP-enhanced thrombosis.

To determine if the SP/mPF4 complex enhances platelet activity, we treated whole blood drawn from K18-hACE2^TG^ and K18-hACE2^TG^/*Tymp^-/-^*mice with COS-7 lysates containing either an SP/mPF4 mixture or mPF4 alone, at a concentration of 50 µg/ml, at 37 °C for 1 hour with gentle rotation. The whole blood was then perfused through a Cellix flow chamber coated with collagen at a shear stress of 60 Dyn/cm^2^. We did not find any difference in platelet adhesion to the collagen coated surface among the tested conditions (**Fig. S7**).

## Discussion

SARS-CoV-2 SP, a structural surface protein that binds to ACE2 to mediate viral entry, may have gained an advantage over its viral protein counterparts in leading to cellular responses and generating a prothrombotic environment even before the virus is significantly amplified. This study, for the first time, validates this hypothesis and provides novel insights into how SARS-CoV-2 SP promotes thrombosis. Several innovative findings in this study include: 1) SARS-CoV-2 SP itself is prothrombotic, by augmenting the intensity and the rate of thrombus formation; 2) targeting TYMP attenuates SP-enhanced thrombosis; 3) SP enhances TYMP expression in human lung epithelial cells. The increase of TYMP also facilitates the expression of SP and PF4, forming a vicious cycle that may worsen disease progression; and 4) SP and TYMP binding enhances the affinity of SP to PF4.

Traditionally, TYMP functions as a DNA salvage pathway enzyme and catalyzes the conversion of thymidine to thymine^55^. Our recent research, utilizing *Tymp*^-/-^ mice, has revealed that TYMP enhances platelet GPVI signaling activation and thus promotes thrombosis^27,56^.

Inhibition of TYMP with tipiracil, an FDA approved drug, significantly inhibits thrombosis without affecting bleeding^28^. As mentioned above, TYMP is significantly increased in the COVID-19 milieu. Using the MGH Olink Proteomics database^29^ and an ROC analysis based on TYMP expression on the day of hospital admission (Day 0), we have found that TYMP serves as a very sensitive and specific marker in diagnosing severe COVID-19 (**Fig. S8**). However, the mechanism underling COVID-19 enhanced TYMP expression as well as the consequent effects or causes remains unclear.

In this study, we show that SP increases TYMP expression in BEAS-2B cells. This effect is mediated by the activation of NF-κB signaling, as evidenced by the simultaneous activation of p65, given that NF-κB is known to augment TYMP expression^50^. While we did not examine the detailed cell populations that respond to SP intraperitoneal injection in vivo, we presume similar responses in K18-hACE2^TG^ mice. This assumption is supported by the evidence of enhanced thrombosis in vivo and increased p-STAT3 expression in platelets isolated from the SP-treated K18-hACE2^TG^ mice. The TYMP-dependent increase in p-STAT3 in platelets aligns with TYMP’s role, as we have reported that TYMP increases serum stimulated STAT3 activation in vascular smooth muscle cells^52^, and both STAT3 and TYMP have been demonstrated to enhance GPVI signaling^27,53^.

Another intriguing finding of our study is that TYMP dose-dependently enhances the affinity between SP and PF4. The pathophysiological significance of the binding between TYMP and SP as well as TYMP-enhanced SP and PF4 binding is not yet clear. The S1 subunit of SP and full-length SP were detected in plasma of acute COVID-19 patients and recently vaccinated patients, with S1 more frequently observed than full-length SP, using an optimized ultra-sensitive single molecule array (Simoa) assay^22,24,25^. S1 was detectable via western blot in our severe COVID-19 patient plasma samples only after immunoprecipitation. Similarly, S1 was detectable in the plasma of mice injected with the SP-containing lysate, but not full-length SP (data not shown), consistent with previous findings in mice vaccinated against full-length SP^57^. A serial dilution of recombinant S1 showed that S1 could not be detected at or below 100 ng/ml via our western blot protocol (Fig. S4). Regardless, a successful immunoprecipitation of S1 (utilizing anti-SP/S1 antibody) and co-IP of PF4 were demonstrated in COVID-19 patient plasma. The implications of a SP/PF4 complex have been suggested in the context of VITT^23,26^, but not in COVID-19 associated thrombosis. PF4 electrostatically enhances the accumulation and aggregation of platelets to endothelial cells, thus facilitating platelet aggregation to form a thrombus^58^. Both PF4 and SP affect coagulation by binding heparan sulfate, which prevents it binding to antithrombin, thus promoting coagulation^19,59,60^. The addition of PF4 to plasma also shortens aPTT^61^. Furthermore, PF4 has recently been shown to activate platelets via a JAK-STAT3-dependent pathway, though concentrations several magnitudes over the physiological, circulating PF4 titer range were used^62^. Additional studies are needed to clarify whether the SP/PF4 complex can amplify PF4 function or synergistically affect coagulation, and whether this complex can trigger some immune response outside of proposed VITT mechanisms and consequently achieve a prothrombotic state.

Puhm et al. suggested tissue factor (TF)- and plasma-dependent coagulation as a potential mechanism of thrombosis in COVID-19^15^. Using the same K18-hACE2^TG^ mice, they found that TF expression was increased starting on day 3 post SARS-CoV-2 infection. Several studies also detected TF activity in the blood of COVID-19 patients^63–65^. As shown in **Fig. S9**, we did not find that SP affects the mRNA expression of TF, FVII, FII, and FXII in either the lungs or the liver on day 3 post-treatment. These findings suggest that either a single injection of SP alone is not sufficient to initiate a change in the mRNA levels of TF, FVII, and other coagulation factors, or the increase of these factors is based on different mechanisms. Several studies also implicated that FVII is increased in the COVID-19 milieu, especially in severe patients^66,67^. In the FeCl_3_-induced thrombosis model, the entire vessel wall of the targeted segment is oxidatively damaged and the endothelial layer is denudated^35,36^. This can lead to TF release in the circulation and generate thrombin through activating factor FVII.

TYMP may potentially affect FVII function. TYMP plays an important role in maintaining mitochondrial DNA (mtDNA) function and a TYMP loss-of-function mutation has been linked to a human mitochondrial disease^55,68^. Integration of mtDNA in the IVS 4 acceptor site of the human factor VII (FVII) gene causes deleterious mutation, resulting in severe FVII deficiency and thereby a bleeding disorder^69^. While FVII lacks an activation peptide, insertion of an activation peptide from other coagulation factors, e.g., FX, dramatically prolonged the half-life of FVII^70^. Additional studies are needed to clarify if TYMP could affect FVII function in the COVID-19 milieu.

The limitation of this study is that we did not use purified full-length SARS-CoV-2 SP. This was due to the unavailability of commercially available full-length SP at the time we initiated this project. Several vendors mentioned that the purified full-length SP is unstable. We thus did not intend to purify full-length SP for this project. However, our home-made protocol generated reproducible data and is considered reliable. Additionally, we employed rigorous controls; all controls (p3.1) and SP lysates were bulk generated using cells from the same passage, and cells originated from the same parent flask.

Taken together, we are the first to have systemically studied the role of SARS-CoV-2 SP in enhancing thrombosis using in vivo mouse models, while providing novel molecular insights using in vitro studies. We found that SP upregulates TYMP expression, and potentially vice versa. Additionally, we demonstrated a binding between full-length SARS-CoV-2 SP and TYMP, with the additional role of TYMP enhancing the formation of the SP/PF4 complex. We conclude that targeting TYMP could be a potential and novel therapeutic strategy for COVID-19-associated thrombosis.

## Acknowledgements

This work is supported by the Marshall University Institute Fund (to WL), the American Heart Association Predoctoral Fellowship (23PRE1018686 to RR), the National Institutes of Health R15HL145573 (to WL) and R15HL164682-01 (to JL), the West Virginia Clinical and Translational Science Institute-Pop-Up COVID-19 Fund (to WL) supported by the National Institute of General Medical Sciences (U54GM104942), as well as the West Virginia IDeA Network of Biomedical Research Excellence WV-INBRE (P20GM103434). This work was also supported in part from NHLBI grants (HL157975-01A1 and HL164016-01A1) to XLZ. The content is solely the responsibility of the authors and does not necessarily represent the official views of the National Institutes of Health.

## Authorship Contributions

Study Design: RR, HY, and WL. Data Collection: RR, YH, AD, EK, FB, YK, KD, AK, QZ, JL, XLZ, and WL. Data Analysis: RR, YK, EK, AK, JL, and WL. Drafting Manuscript: RR and WL. Critical Revisions: HY, YK, JL, and XLZ.

