## Supplemental Methods and Materials for "Thymidine Phosphorylase Mediates SARS-CoV-2 Spike Protein Enhanced Thrombosis in K18-hACE2^TG^ Mice"

**Running title:** TYMP mediates SARS-CoV-2 Spike-enhanced Thrombosis

Corresponding author:
Wei Li, MD, PhD, FAHA
Department of Biomedical Sciences, BBSC 241G
Joan C. Edwards School of Medicine
Marshall University
One John Marshall Drive, Huntington,
West Virginia 25755-9310, USA.


### Methods

#### Plasmids

Plasmids pcDNA3.1-SARS-CoV-2-S-RBD-sfGFP (Item# 141184), pcDNA3.1-SARS2-Spike (Item# 145032), and PF4-bio-His (Item# 53411) were purchased from Addgene under a Material Transfer Agreement. PF4-bio-His was used as DNA template and human PF4 was amplified by PCR and cloned into pCDNA6B-His with the cloning sites HindIII and EcoRI. The newly constructed plasmid pCDNA6B-hPF4-His was used in this project. Mouse PF4 plasmid, pMD-mPF4, was purchased from Sino Biological. Plasmids pcDNA6B-hTYMP or pcDNA3.1-hTYMP have been previously generated in the Li lab and routinely used in different projects.

#### Antibodies

| Protein Target | Manufacturer | Catalog # | Use |
|---|---|---|---|
| SP S1 subunit | Cell Signaling Technology, USA | 99423 | Western blot; immunocytofluorescence; immunoprecipitation |
| SP S1 subunit (RBD) | Cell Signaling Technology, USA | 69323 | Western blot; immunoprecipitation |
| SP NTD | Cell Signaling Technology, USA | 56996 | Immunoprecipitation |
| p-STAT3 | Cell Signaling Technology, USA | 9145 | Western blot |
| T-STAT3 | BD Biosciences, USA | 610190 | Western blot; immunoprecipitation |
| p-JAK2 | Cell Signaling Technology, USA | 3771 | Western blot |
| T-JAK2 | Cell Signaling Technology, USA | 3229 | Western blot |
| His | Cell Signaling Technology, USA | 12698 | Western blot; immunocytofluorescence |
| His | Thermo Fisher Scientific, USA | MA1-21315 | Immunoprecipitation |
| α-Tubulin | GenScript Biotech, USA | A01490 | Western blot |
| GAPDH | Cell Signaling Technology, USA | 51332 | Western blot |
| TYMP | Abcam | ab180783 | Western blot |
| TYMP | Santa Cruz Biotechnology, USA | Sc-47702 | Immunoprecipitation; western blot |
| Mouse Gamma Globulin | Thermo Fisher Scientific, USA | 31878 | Immunoprecipitation |
| Rabbit Gamma Globulin | Thermo Fisher Scientific, USA | 31877 | Immunoprecipitation |

| | | | |
|---|---|---|---|
| PF4 | Santa Cruz Biotechnology, USA | sc-398979 | Western blot |
| p-p65 | Cell Signaling Technology, USA | 3033 | Western blot |
| T-p65 | Cell Signaling Technology, USA | 6956 | Western blot |
| PSP-C | Abcam | ab90716 | Immunofluorescence |
| mPF4 | R&D Systems, USA | AF-595 | Immunoprecipitation |
| Myc | Sigma-Aldrich, USA | M4439 | Immunoprecipitation |
| Anti-mouse IgG, HRP-linked | Cell Signaling Technology, USA | 7076 | Western blot |
| Anti-rabbit IgG, HRP-linked Antibody | Cell Signaling Technology, USA | 7074 | Western blot |

**SARS-CoV-2 Spike Protein Generation**

Using the FuGENE® 6 Transfection Reagent protocol (Cat# E2691, Promega, USA), we transiently transfected pcDNA3.1-SARS-CoV-2-S-RBD-sfGFP and pcDNA3.1-SARS2-Spike into COS-7 cells to generate homemade crude SP- and RBD-containing cell lysates. COS-7 cells were seeded at 1.5E+06 cells/10 cm cell culture dish and cultured in Dulbecco's Modified Eagle's Medium (DMEM) with 10% fetal bovine serum (FBS, S11550H, ATLANTA Biologicals) and 1x antibiotic-antimycotic (Cat# 15240062, Thermo Fisher Scientific, USA). A total of 6 µg plasmids was used for each transfection. The cells were allowed to grow for 36 hours, were washed three times with cold phosphate-buffered saline (PBS) to remove FBS and cell residue, and then were scraped into 500 µL PBS without adding any inhibitors. The cells were sonicated three times in ice, and centrifugated at 12,000 g for 10 min at 4 ºC. The supernatant was transferred into a new clean tube and protein concentration was measured using the Bradford protein assay (Cat# 5000201, Bio-Rad, USA). The presence and relative concentration of SARS-CoV-2 SP or RBD was confirmed by a western blot using the recombinant SP S1 subunit (Cat# P1532, BioVision, USA) as a control **(Fig. S1).** COS-7 cells transfected with empty plasmid pcDNA3.1 were used as a control (p3.1).

**Animals**

K18-hACE2$^{TG}$ mice were purchased from The Jackson Laboratory (USA). K18-hACE2$^{TG}$ mice were crossed with *Tymp*$^{-/-}$ mice to generate K18-hACE2$^{TG}$/*Tymp*$^{-/-}$ mice. *Tymp*$^{+/-}$ mice have been previously back-crossed with C57BL/6J mice for ten generations[1]. Pups were tagged at the time of weaning and genotyped based on the protocol provided by the Jackson Laboratory. Mice aged 8-16 weeks were used for experiments. Mice were intraperitoneally injected with 500 µg SP-containing or p3.1 COS-7 cell lysate and 72 hours later, mice were subjected to the thrombosis assay in vivo, or were harvested for platelets and plasma for platelet aggregometry, measuring activated partial thromboplastin time (aPTT), or platelet biochemistry assays.

**Blood Collection**

Mice were anesthetized with ketamine/xylazine (100/10 mg/kg) by intraperitoneal injection. Whole blood was drawn via the abdominal inferior vena cava puncture using 0.109 M sodium citrate as an anti-coagulant at 1:9 ratio by volume.

#### Platelet Aggregometry

Modified Tyrode's HEPES buffer was added to the whole blood at a ratio of 7:10 volume, and centrifuged at room temperature, 100 g for 10 min. The supernatant, namely platelet-rich plasma (PRP) was transferred into a new fresh tube. The remaining cellular portion was further centrifuged at high speed (>10,000 rpm) for 15 seconds to receive platelet-poor plasma (PPP). PPP was also transferred into a new tube. Platelets were counted with a hemocytometer and were adjusted to 2.5E+08 cells/mL with corresponding PPP and 300 µL of the platelet-adjusted PRP was used in the aggregation assay. Samples were stirred at 1,000 RPM at 37 ºC. Calcium chloride/magnesium chloride ($CaCl_2$/$MgCl_2$) was added to PRP to a final concentration of 1 mM immediately before adding collagen (Cat# 385, Chrono-log, USA) or adenosine diphosphate (ADP) (Cat# 384, Chrono-log, USA). A Model 700 aggregometer (Chrono-log, USA) paired with Aggrolink8 version 1.29.29–31 on Windows XP Professional was used to monitor light transmission over time. Aggregation was quantified with 100% aggregation corresponding to 100% light transmission[2,3].

#### Activated Partial Thromboplastin Time (aPTT)

Blood was drawn as mentioned above and PPP was separated by centrifugation at room temperature, 1,000 g for 6 min. The top two-thirds of the supernatant was transferred into a new tube. 150 µL of PPP was used per assay. Samples were stirred at 1,000 RPM at 37 ºC. 150 µl of $CaCl_2$ (Cat# 21-405, R2 Diagnostics, USA) was added to the PPP followed by 150 µl phospholin ES reagent (Cat# 21-405, R2 Diagnostics, USA). Light transmission was monitored over time via the aggregometer and software described above. aPTT was obtained by determining the time required for a light transmission change after the addition of phospholin ES.

#### Washed Platelet Workup for Protein Expression Studies

Blood cells were resuspended with sodium citrate in PBS (1:9 in vol/vol) in the fractionated whole blood remaining after collecting PPP from the aPTT procedure. PRP was separated as mentioned in the Platelet Aggregation section and platelets were pelleted and resuspended with sodium citrate in PBS with prostaglandin E1 (PGE1) (Cat# P5515, Sigma-Aldrich, USA) (500 ng/µL). The platelets were washed once more with sodium citrate in PBS with PGE1 and were resuspended in 100 µl RIPA with Halt protease/phosphatase inhibitors (Cat# 1861280, Thermo Fisher Scientific, USA). Standard western blotting protocol was performed. Protein signals were quantified using ImageJ software.

#### Ferric Chloride ($FeCl_3$) Injury-Induced Murine Carotid Artery Thrombosis Model

The $FeCl_3$ injury-induced carotid artery thrombosis model has been previously described in extensive detail[4,5]. Still images were taken at minute intervals for the first 6 min after injury and, via ImageJ software (NIH, USA), integrated density, area, and mean grey value of whole images were measured. A background measurement was taken outside of the vessel. Fluorescence intensity was calculated by (integrated density) - (the whole image area X mean value of background measurement).

#### Treating BEAS-2B cells with SARS-CoV-2 SP for Western Blotting

BEAS-2B cells, a human bronchial epithelial cell line, were seeded at 5.0E+05 cells/well in six-well plates pre-coated one night before with a mixture of 0.01 mg/mL fibronectin, 0.03 mg/mL bovine collagen type I, and 0.01 mg/mL bovine serum albumin dissolved in BEBM medium based on ATCC protocol. The complete medium for this cell line (BEBM) is Airway Epithelial Cell Basal Medium (PCS-300-030) and Bronchial Epithelial Cell Growth Kit (PCS-300-040). Cells were allowed to adhere to the plate bottom for six hours, and then treated with 100 µg (50 µg/ml) SARS-CoV-2 SP- or RBD-containing or control COS-7 cell lysate for 24 hours. Cells were washed with cold PBS for three times and then harvested in 100 µl RIPA with Halt protease/phosphatase inhibitors and standard western blotting protocol was performed.

#### TYMP siRNA Transfection in BEAS-2B cells

The Lipofectamine™ 3000 Transfection Reagent protocol (Cat# L3000-015, Thermo Fisher Scientific, USA) was used for siRNA transfection. BEAS-2B cells at a concentration of 1E+05 cells/well in six-well plates were transfected with 5 nM TYMP siRNA (Cat# 4392420, Thermo Fisher Scientific, USA), or scramble siRNA (Cat# 4390843, Thermo Fisher Scientific, USA) as negative control, six hours after seeding. Cells were allowed to grow for an additional 48 hours and then treated with 100 µg SARS-CoV-2 SP-containing COS-7 cell lysate for 24 hours. Cells were washed with cold PBS and then lysed in RIPA buffer and used for western blot assay.

#### Western blotting

Protein samples were mixed with 2x Laemmli Sample Buffer (Cat# 1610737, Bio-Rad, USA), heated at 95 °C for 5 min, and then dissolved into sodium dodecyl sulfate–polyacrylamide gel electrophoresis (SDS-PAGE). The separated proteins were transferred onto polyvinylidene difluoride (PVDF) membrane in transfer buffer (25 mM Tris, 192 mM glycine, 20% v/v methanol). The PVDF membrane was blocked with 5% milk in PBS for 1 hour at room temperature with gentle rocking and then incubated with the primary antibody diluted in 5% milk in PBS in a concentration suggested by the manufacturer for overnight at 4 °C. The membrane was washed with tris-buffered saline containing 0.1% Tween 20 (TBS-T) for three 15 min intervals, incubated with the secondary antibody at RT for 1 hour, washed with TBS-T for three 15 min intervals once more, and then developed with ECL kit(s) (Cat# 34076, 34580, A38554 Thermo Fisher Scientific).

#### Co-immunoprecipitation (IP) with platelet factor 4, SP, and TYMP

Using Lipofectamine™ 3000, we transfected COS-7 cells seeded at 1.5E+06 cells/10 cm culture dish with combinations of platelet factor 4 (PF4)-, SARS-CoV-2 SP-, and Tymp-containing plasmids to test potential binding among the expressed proteins. A total of 6 µg plasmids was transfected at each condition. Transfected cell lysates were collected in immunoprecipitation (IP) lysis buffer (25 mM Tris-HCl pH 8, 150 mM NaCl, 1% NP-40, 1 mM EDTA, 5% glycerol) containing Halt protease/phosphatase inhibitors. One milligram of the protein lysates was pre-cleaned with 300 µg Dynabeads™ Protein G for Immunoprecipitation (Cat# 10004D, Thermo Fisher Scientific, USA) and then were incubated overnight with IP antibody in a 1.5 mL tube, with gentle rotation at 4 ºC. The resulting mixtures were then incubated with 1.5 mg Dynabeads™ for 2 hours at 4 ºC. The supernatant was decanted and the Dynabeads™ were washed in cold PBS 3x for 5 min at 4 ºC and eluted in 30 µl 2x Laemmli Sample Buffer (Cat# 1610737, Bio-Rad, USA) at 95 ºC for 5 min. The resulting eluates were analyzed via western blotting alongside their respective inputs.

**Immunocytochemistry with platelet factor 4 and SP**

COS-7 cells seeded at 1.5E+05 cells/Nunc™ glass bottom dish were transfected with PF4- and SP-containing plasmids (3 µg for each plasmid, total 6 µg) using Lipofectamine™ 3000. Cells were washed with cold PBS 2x, fixed in 4% paraformaldehyde in PBS for 10 min at room temperature, and permeabilized with Triton X-100, followed by washing in PBS 3x for 5 min each. Blocking was performed with 5% milk in PBS for 1 hour at room temperature followed by simultaneous incubation with anti-His and anti-SP overnight at 4 ºC in the dark. Cells were washed in PBS 3x for 5 min each and were incubated simultaneously with Alexa Fluor® 488 (Cat# A-11008, Thermo Fisher Scientific, USA) against anti-SP and Alexa Fluor® 568 (Cat# A-11004, Thermo Fisher Scientific, USA) against anti-His for 1 hour at room temperature in the dark. Cells were washed 3x with PBS for 5 min each and were mounted with VECTASHIELD® Antifade Mounting Medium with DAPI (Cat# H-1800, VectorLabs, USA). A Leica DMi8 inverted microscope (Leica, Germany) and CaptaVision+™ Software (Accu-scope, USA) were used to visualize co-immunofluorescence.

**Blue Native Polyacrylamide Gel Electrophoresis for Identifying SP and PF4 Interaction**

The sample preparation and analysis for blue native polyacrylamide gel electrophoresis (BN-PAGE) followed a previous publication[6] that was adopted from the protocol described by Schägger and Wittig et al[7-9]. COS-7 cells co-transfected with plasmids encoding SP and PF4 were lysed with 1X BN-PAGE sample buffer (with 2% DDM, n-dodecyl-β-D-maltoside and Halt protease/phosphatase inhibitors) and cleared by centrifuge at 20,000 g for 30 min at 4°C and stored at -80°C until use. Immediately before sample loading, the NativePAGE 5% G-250 sample additive was added to samples (at 1/4th of detergent DDM concentration) and mixed. Electrophoresis was conducted according to the manufacturer's instructions. Proteins dissolved in the gel were transferred onto PVDF membrane and followed standard immunoblotting procedures using antibodies against SP and His-tagged PF4.

**Serial Dilution of S1 SP**

Recombinant SP S1 subunit (Cat# P1532, BioVision, USA) was serially diluted from 1 μg/ml to 1 pg/ml, and then subjected to a western blot assay. Immunoblotting followed standard western blotting protocol and was performed with anti-S1 (Cat# 99423S, Cell Signaling Technology, USA) at 1:1000 at 4 °C overnight and HRP-linked anti-IgG (Cat# 7074, Cell Signaling Technology, USA) at 1:2000 at RT for 1 hour. Chemiluminescent signal was detected using ECL kit (Cat# 34076, Thermo Fisher Scientific) and 5 minutes of exposure was allowed before film processing.

#### qPCR of Murine Lungs and Liver

Mice were exsanguinated via inferior vena cava and cold PBS was perfused through the left ventricle. The liver and lungs were harvested and snap frozen in liquid nitrogen. RNA was extracted via the Qiagen RNeasy Mini Kit (Cat# 73404, Qiagen, Germany). cDNA was synthesized using the SuperScript™ VILO™ cDNA Synthesis Kit (Cat# 11754-050, Thermo Fisher Scientific, USA), and PowerUp™ SYBR™ Green Master Mix (Cat# A25742, Thermo Fisher Scientific, USA) was used as a reporter for qPCR. StepOnePlus Real-Time PCR System (Thermo Fisher Scientific, USA) with StepOne Software v2.3 (Thermo Fisher Scientific, USA) were used to ascertain relative cDNA concentrations which were analyzed using the $2^{-\Delta\Delta CT}$ method-based calculation.

**Primers**

| Gene target | Forward Primer | Reverse Primer |
|---|---|---|
| FII (prothrombin) | 5'-GCC AAG ACC CTG AGC AAG TA-3' | 5'-TCG TCC ACA TCA TAG TTC TCC T-3' |
| FIII (Tissue factor) | 5'-AGT TCA TGG AGA CGG AGA CC-3' | 5'-TTG CCA AAG ACT TGC CGC-3' |
| FVII (Factor VII) | 5'-TGC CAG GAT CAT CTC AAG TCT-3' | 5'-AGC TAC AGG TAC GCT TGG TC-3' |
| FXII (Factor XII) | 5'-CAG GCT GGA ACT ACG CAA TC-3' | 5'-CAG GGA CAA CTA GAG GTG CA-3' |

#### AlphaFold2-Based Binding Prediction

AlphaFold2, an advanced deep-learning model for protein structure prediction, predicted the binding interface between TYMP or PF4 and SP through the ChimeraX interface. The complete TYMP sequence served as the basis for the prediction. Regarding SP, we employed four distinct domain sequences due to the considerable size of the full-length protein, making binding prediction challenging. The final binding model was established by substituting each protein structure retrieved from the Protein Data Bank (PDB) into the AlphaFold2 prediction model, implemented through the Matchmaker function in ChimeraX. To ensure their representativeness and relevance for the binding prediction analysis, TYMP and SP structures were selected from PDB, closely related by sequence alignment. Visualizations of the predicted binding interfaces were created using ChimeraX, a molecular visualization tool[10,11]. To analyze potential binding interfaces, we meticulously examined AlphaFold2 predictions, aiming to identify potential binding interactions between TYMP and SP. The Structure Analysis tools in ChimeraX were then utilized to measure distances between potentially interacting side chains.

**Collagen-Mediated Platelet Adhesion Assay**

A Vena8 Fluoro+ 8-channel microfluidic flow chamber (Cellix, Ireland) was coated with 100 µg/ml collagen (Cat# 385, Chrono-log, USA) and was incubated at 4 ºC overnight. Whole blood collected from untreated mice was incubated with a PF4-containing and a dual PF4, SP-containing COS-7 cells lysate (50 µg/ml) for 1 hour at 37 ºC. An ExiGo pump was set at a constant flow rate of 60 µl/min and SmartFlo software (Cellix, Ireland) designed for the Apple iPad (Apple, USA) operated the pump. Rhodamine 6G was added into the treated blood in a final concentration of 25 µg/mL to staining platelets. After 3 minutes of running of the blood sample, PBS was perfused in the same flow rate for additional 3 minutes and then still images were taken at predesignated points on each channel with a Leica DMi8 inverted microscope and CaptaVision+™ Software. The fluorescence area was quantified via ImageJ software using brightness as a color threshold until the background area outside of the chamber showed no signal.

**Human Plasma Preparation**

Ethylenediaminetetraacetic acid (EDTA)-anticoagulated blood samples were obtained on admission from patients with severe and critical COVID-19. The whole blood was centrifugated at 1500 rpm for 15 min. Platelet-poor plasma was collected and stored in aliquots at -80 °C until usage.
