## Supplemental Figures for "Thymidine Phosphorylase Mediates SARS-CoV-2 Spike Protein Enhanced Thrombosis in K18-hACE2^TG^ Mice"

### Supplemental Figure 1

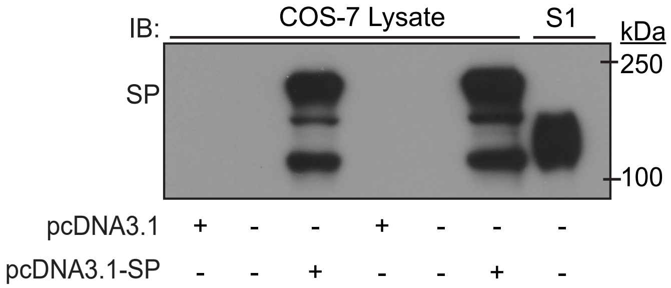


**Supplemental Figure 1. SARS-CoV-2 spike protein is reproducibly expressed in COS-7 cells transfected with SP-containing plasmids.** COS-7 cells were transiently transfected with plasmids encoding full length SARS-CoV-2 spike protein (SP), or its receptor binding domain (RBD), or the empty backbone vector pCDNA3.1 using FuGENE 6. Cell lysates were prepared 36 hours later and 30 µg total proteins was analyzed by western blot. 1 µg of SP S1 subunit purchased from BioVision was run in the same gel as control.

**Supplemental Figure 2**

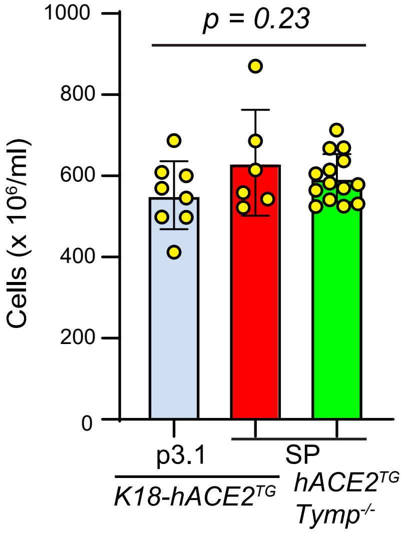


**Supplemental Figure 2. SARS-CoV-2 spike protein treatment does not affect platelet titers in K18-hACE2$^{TG}$ mice.** Whole blood was drawn from mice with 0.109M sodium citrate (1:9) as an anticoagulant and was immediately analyzed with a Drew Scientific 950FS Hemavet. Data were analyzed with one-way ANOVA. p3.1: lysate from COS-7 cells transiently transfected with pCDNA3.1 as control. SP: lysate from COS-7 cells transiently transfected with pCDNA3.1-SARS-CoV-2-SP.

#### Supplemental Figure 3

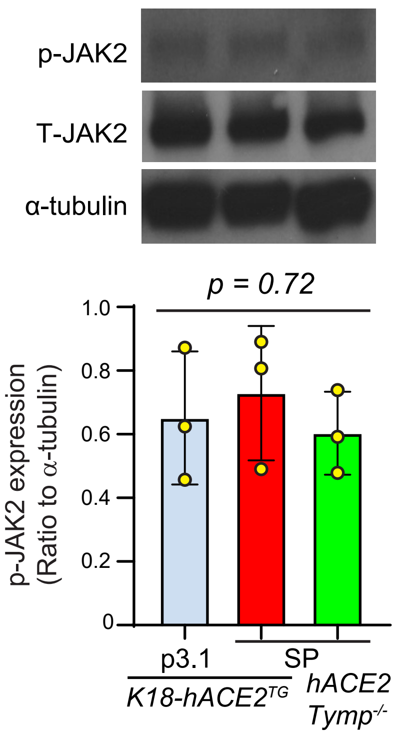


**Supplemental Figure 3. SARS-CoV-2 spike protein does not affect Janus kinase 2 activation in K18-hACE2$^{TG}$ platelets.** The expression of phosphorylated (p) and total (T) JAK2 were probed via western blot in murine platelets harvested from SP- or control-treated mice. Alpha-tubulin was blotted as a loading control. Data were analyzed by one-way ANOVA.

#### Supplemental Figure 4

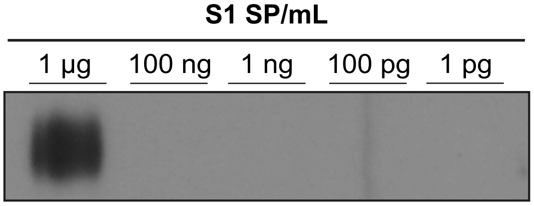


**Supplemental Figure 4. Titration of the sensitivity of western blot on detecting SARS-CoV-2 SP expression.** Recombinant SP S1 subunit (Cat# P1532, BioVision) was serially diluted from 1 μg/ml to 1 pg/ml and detected via western blot using the anti-S1 antibody (Cat# 99423, Cell Signaling Technology) at a concentration of 1:1000. A 1:2000 concentration was used for the secondary antibody (Cat# 7074, Cell Signaling Technology). Signal was detected using ECL kit (Cat# 34076, Thermo Fisher Scientific).

#### Supplemental Figure 5

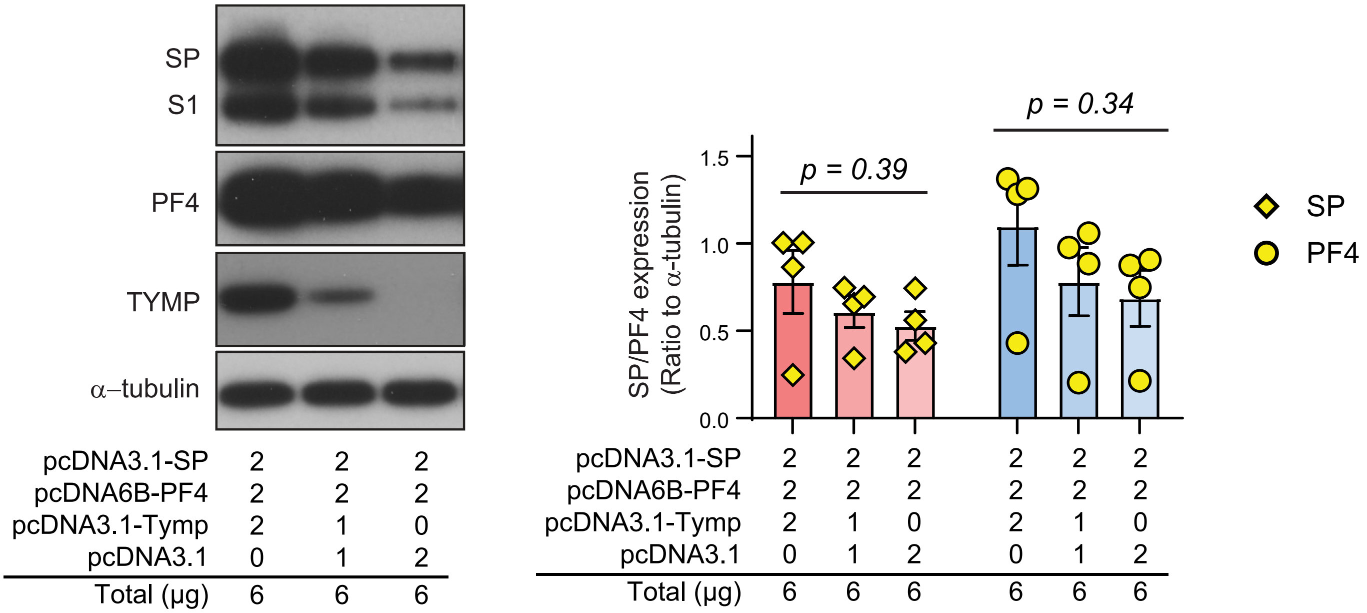


**Supplemental Figure 5. TYMP overexpression tends to increase the expression of SARS-CoV-2 SP and platelet factor 4 (PF4).** COS-7 cells were co-transfected with plasmids encoding SARS-CoV-2 SP, human PF4, and human TYMP. SP- and PF4-plasmids were maintained at a ratio of 1:1 in mass while varying TYMP-plasmid concentrations. The resulting cell lysates were prepared for western blot to assess the expression of SP, PF4, TYMP, and α-tubulin. Expression data were analyzed with one-way ANOVA.

#### Supplemental Figure 6

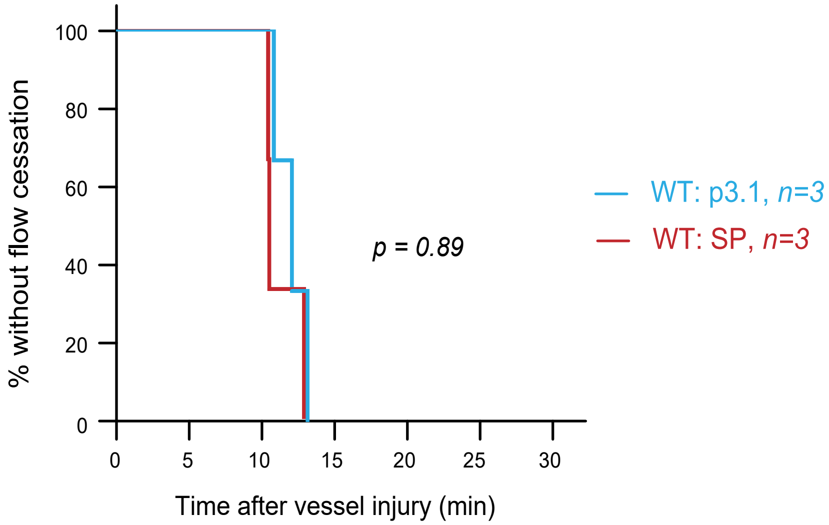


**Supplemental Figure 6. SARS-CoV-2 spike protein does not enhance thrombosis in C57BL/6 mice.** C57BL/6J mice were intraperitoneally injected with 500 μg of SARS-CoV-2 spike protein-containing or control (p3.1) COS-7 cell lysate. The mice were subjected to the 7.5% FeCl3-induced carotid artery thrombosis model 72 hours after SP or control treatment. Data are shown as a survival plot of time to flow cessation after vessel injury. Groups were compared using the logrank test.

#### Supplemental Figure 7

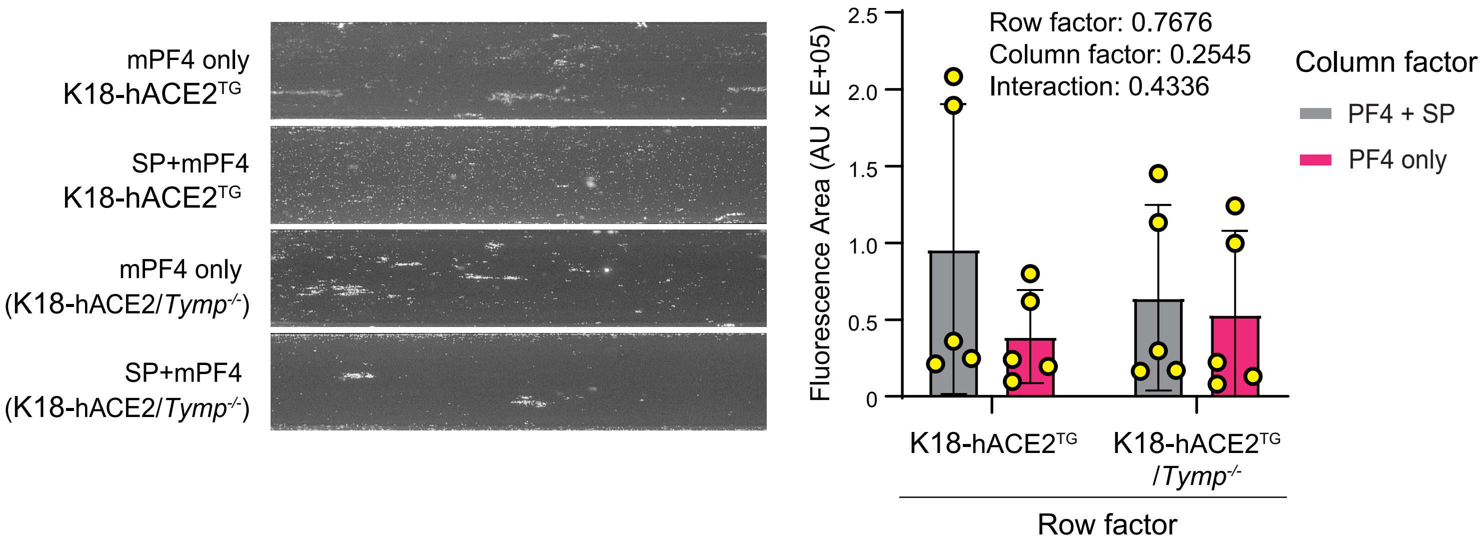


**Supplemental Figure 7. The complex of SARS-CoV-2 SP and PF4 does not significantly promote platelet adhesion to collagen-coated surface.** Whole blood from K18-hACE2$^{TG}$ and K18-hACE2TG/*Tymp*$^{-/-}$ mice was incubated with SP/murine PF4 (mPF4) or mPF4-only COS-7 lysate (50 ug/mL) at 37 $^{o}$C for 1 hour. The treated blood was then perfused through a collagen-coated Cellix flow chamber at a shear force of 60 Dyn/cm² for 3 min. The chambers were washed with PBS at the same flow rate for another 3 minutes and then images were taken at predesignated marker positions 2 to 6. Representative images are shown on the left panel. Fluorescence areas, representating areas covered by platelets, were quantified by ImageJ and were analyzed via a two-way ANOVA.

**Supplemental Figure 8**

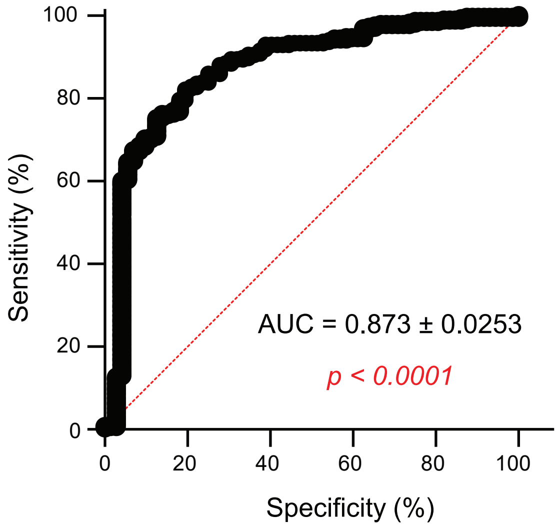


**Supplemental Figure 8. Diagnostic Value of Thymidine Phosphorylase (TYMP) in COVID-19.** Data on TYMP plasma levels of COVID-19 patients on day of admission were extracted from the MGH Olink Proteomics database as previously mentioned (Li and Yue, Front Med (Lausanne). 2021 Mar 22:8:653773). Seventy-two COVID-19 negative patients were used as controls, while 285 COVID-19 positive individuals were considered patients and used for Receiver Operating Characteristic (ROC) analysis. The area under the curve (AUC) is 0.8721, suggesting that TYMP is a highly sensitive and specific marker for diagnosing COVID-19.

#### Supplemental Figure 9

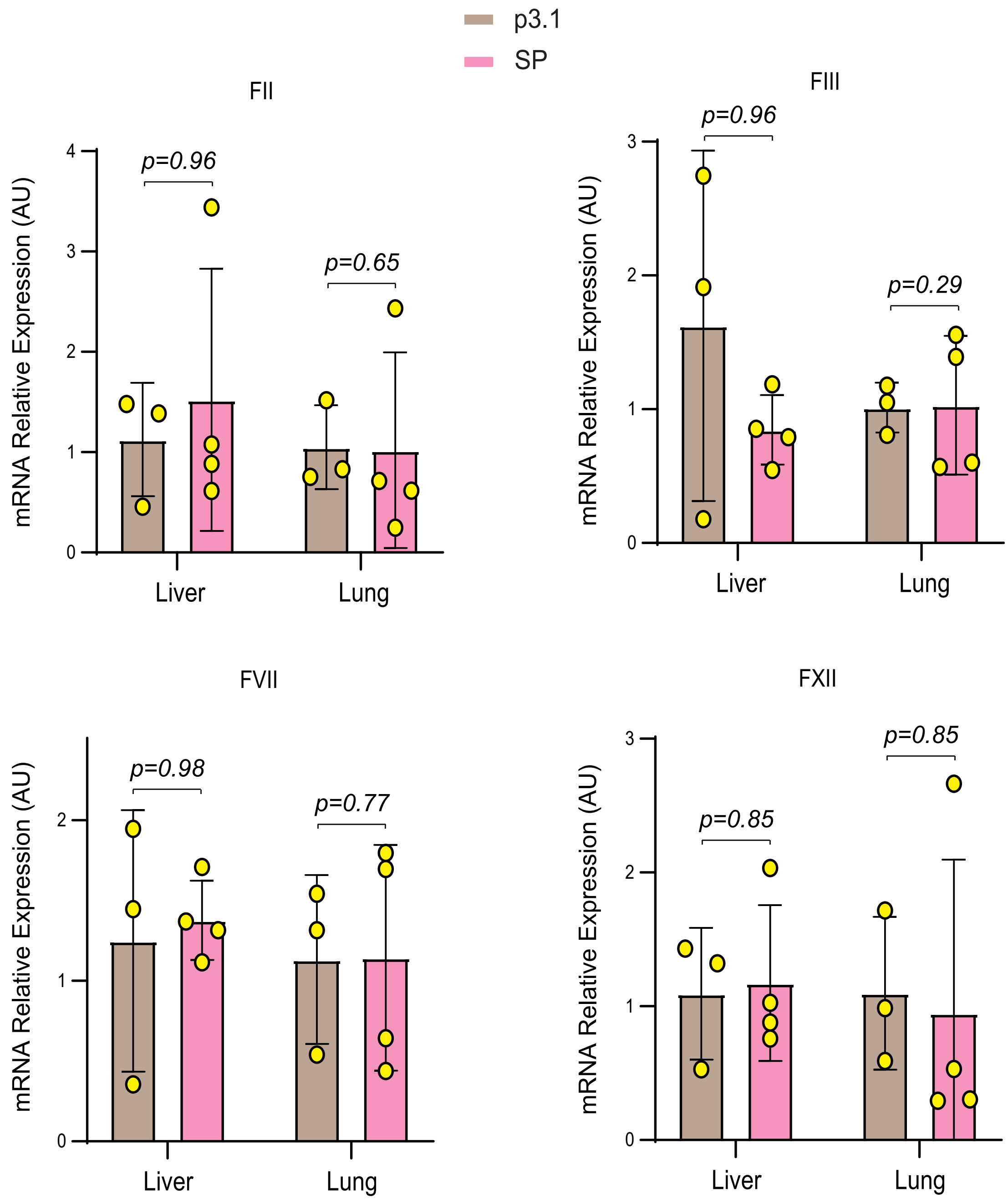


**Supplemental Figure 9. Effect of SARS-COV-2 SP on the expresion of several coagulation factors.** RNA was extracted from the lungs and livers of K18-hACE2$^{TG}$ mice treated with p3.1- or SP-containing COS-7 lysate (500 µg/mouse) using the Qiagen RNeasy Mini Kit. cDNA was synthesized using the Super-Script™ VILO™ cDNA Synthesis Kit. Quantative PCR was conducted to analyze the expression of prothrombin (FII), tissue factor (FIII), factor VII (FVII), and factor XII (FXII) using the SYBR Green qPCR Master Mix kit. 18S ribosomal RNA was used as a housekeeping control gene. The threshold cycle values automatically generated by the qPCR system were used for the $2^{-\Delta\Delta CT}$ method-based calculation of each gene expression. Stuent's *t*-test was used.
